# A transcriptional control model for *doublesex*-dependent sex differentiation in *Nasonia* wasps

**DOI:** 10.1101/2025.02.03.636189

**Authors:** Julien Rougeot, Filippo Guerra, Eveline C. Verhulst

**Author notes:** These authors contributed equally to this work (Julien Rougeot, Filippo Guerra).

## Abstract

The transcription factor Doublesex (DSX) orchestrates insect sex differentiation. DSX affects *Drosophila melanogaster* male and female transcriptome, yet how DSX regulates gene expression in other species is poorly understood. We investigated sex-biased gene expression during the development of the parasitoid wasp *Nasonia vitripennis*, finding that more than three-quarters of its genes are sex-biased in at least one developmental point. Next, we transiently knocked down *dsx* expression to infer its role in sex-specific transcriptome regulation, revealing thousands of affected genes in males and a more subtle effect in females. Finally, we performed an *in vitro* DNA-protein interaction assay to identify DSX binding sites on the genome and primary DSX target genes. By integrating these three datasets, we defined DSX’s regulatory function for all genes in *N. vitripennis*, revealing that DSX acts mainly in males as both an activator and a repressor. This male-centric model for DSX-mediated regulation is likely to apply to many other insect species.

## 1 Introduction

In insects, sexual differentiation is established during early development by the functionally conserved transcription factor Doublesex (DSX) (reviewed in Bachtrog et al., 2014; Salz, 2011; Verhulst et al., 2010). In most species, the *dsx* primary transcript undergoes sex-specific splicing (Chikami et al., 2022; Wexler et al., 2019) that yields male (DSX^M^) and female (DSX^F^) specific protein isoforms. These isoforms contain a highly conserved N-terminal DNA binding domain common to both male and female isoforms and an isoform-specific C-terminal oligomerisation domain. The DNA-binding domain recognises a DNA sequence that is partially conserved among mammals (Murphy et al., 2010; Raymond et al., 2000), roundworms (Yi & Zarkower, 1999) and insects (Burtis et al., 1991; Luo et al., 2011; Suzuki et al., 2003), while the C-terminal domain is thought to evolve fast and is responsible for the sex-specific regulation of DSX target genes (Shukla & Nagaraju, 2010; Verhulst & van de Zande, 2015).

Genomic studies on *Drosophila melanogaster* revealed thousands of sex-biased genes that are potential targets of DSX (Arbeitman et al., 2004; Assis et al., 2012; Chang et al., 2011; Fujii & Amrein, 2002; Goldman & Arbeitman, 2007; Jin et al., 2001; Ranz et al., 2003; White et al., 2002; Whittle & Extavour, 2019). Moreover, direct DSX targets were defined by genome-wide analyses of DSX occupancy in different tissues and both sexes (Clough et al., 2014; Luo et al., 2011). By coupling transcriptomic analyses with functional studies, different modes of gene expression regulation by DSX were discovered: early studies suggested an opposing effect in males and females (Belote et al., 1985; Burtis et al., 1991), while current models predict that DSX can act as a repressor or an activator, in one or both sexes simultaneously (Arbeitman et al., 2016; Goldman & Arbeitman, 2007). Identification of sex-biased gene expression was then pursued in other *D. melanogaster* strains (Arbeitman et al., 2016), Drosophilidae species (Assis et al., 2012; Khodursky et al., 2020; Ranz et al., 2003; Williams et al., 2008), and other insect species (Ledón-Rettig et al., 2017; Wang et al., 2020). These studies showed that DSX regulates thousands of genes in all examined species, in both males and females.

However, these studies rely on species that belong to the derived Aparaglossata taxon, a clade including all Holometabola except the Hymenoptera. Chickami et al. (2022) proposed that the functionalisation of DSX in females arises from the evolution of a disordered region at the DSX^F^ C-terminus specific to the Aparaglossata, whilst in most other species DSX is expected to control the male transcriptome. This suggests that the “canonical” model for DSX-dependent gene regulation developed in *D. melanogaster* might be a novelty and not a general feature of insect sex determination. Therefore, we sought to expand this model by studying the developmental dynamics and different DSX target gene regulation modes in a Hymenoptera model species, the parasitoid wasp *Nasonia vitripennis*.

*N. vitripennis* exhibits multiple sexual dimorphic traits, such as dark pigmentation on the femora and antennae in females and short and narrow wings in males (Darling & Werren, 1990). *N. vitripennis* has a high-quality and fully annotated genome (Dalla Benetta et al., 2020; Werren et al., 2010), and the use of double-stranded RNA (dsRNA) to induce transient, systemic gene knockdown via RNA interference (RNAi) is well established (Lynch, 2015; Lynch & Desplan, 2006; Wang et al., 2022a, Wang et al., 2022b). The sex determination pathway is well described in *N. vitripennis,* which terminates with *dsx* primary transcript being spliced into one female and three male isoforms (Wang et al., 2022a). All isoforms share the canonical N-terminal DNA-binding domain, of which the DNA-binding sequence is unknown. A microarray approach at different developmental stages identified 43% of all *N. vitripennis* genes as having sex-biased expression (Rago et al., 2020). The number of sex-biased genes progressively increases from a few hundred in embryos to more than 6,000 genes in whole adults (Rago et al., 2020). An RNA-seq approach comparing *N. vitripennis* and *N. giraulti* whole adult transcriptomes identified that 75% of all genes have sex-biased expression in either species; with such a big number of sex-biased genes, transcriptomes from the same sex from different species are more similar than those from different sexes from the same species (Wang et al., 2015). Comparative analysis of germline tissues from male and female *N. vitripennis* identified genes expressed uniquely in these tissues (Ferree et al., 2015). Nevertheless, a model for how *dsx* regulates sex-biased gene expression in *N. vitripennis* has yet to be presented.

We use the *Nasonia* model system to examine transcriptome-wide changes in sex-biased gene expression following transient silencing of *dsx* across development and in different tissues. We identify the DSX-binding motif using DNA affinity purification and sequencing (DAP-seq) and reveal hundreds of possible direct DSX-target genes. By synergistically combining the RNA- and DAP-seq datasets, we define a model for DSX regulation of its primary targets.

## 2 Materials and Methods

### 2.1 Insect rearing

For all experiments, *Nasonia vitripennis* lab strain AsymCX was reared on *Calliphora sp.* pupae under a 16/8 hours light/dark cycle at 25°C. *N. vitripennis* AsymCX was established by researchers at the Evolutionary Genetics Department (University of Groningen, The Netherlands) and has been reared for more than 100 generations at the entomology department at Wageningen University (NL).

### 2.2 Transcriptome analysis

We performed transcriptome analysis of male and female *N. vitripennis* at five different life stages after subjecting them to dsRNA injections against either *dsx* or the exogenous gene GFP. This experimental setup allowed us to define the genes that are sex-biased and those that are under DSX regulatory control. In the next paragraphs, we detail our experimental approach, which is schematised in **Figure 1A**.

**Figure 1:**
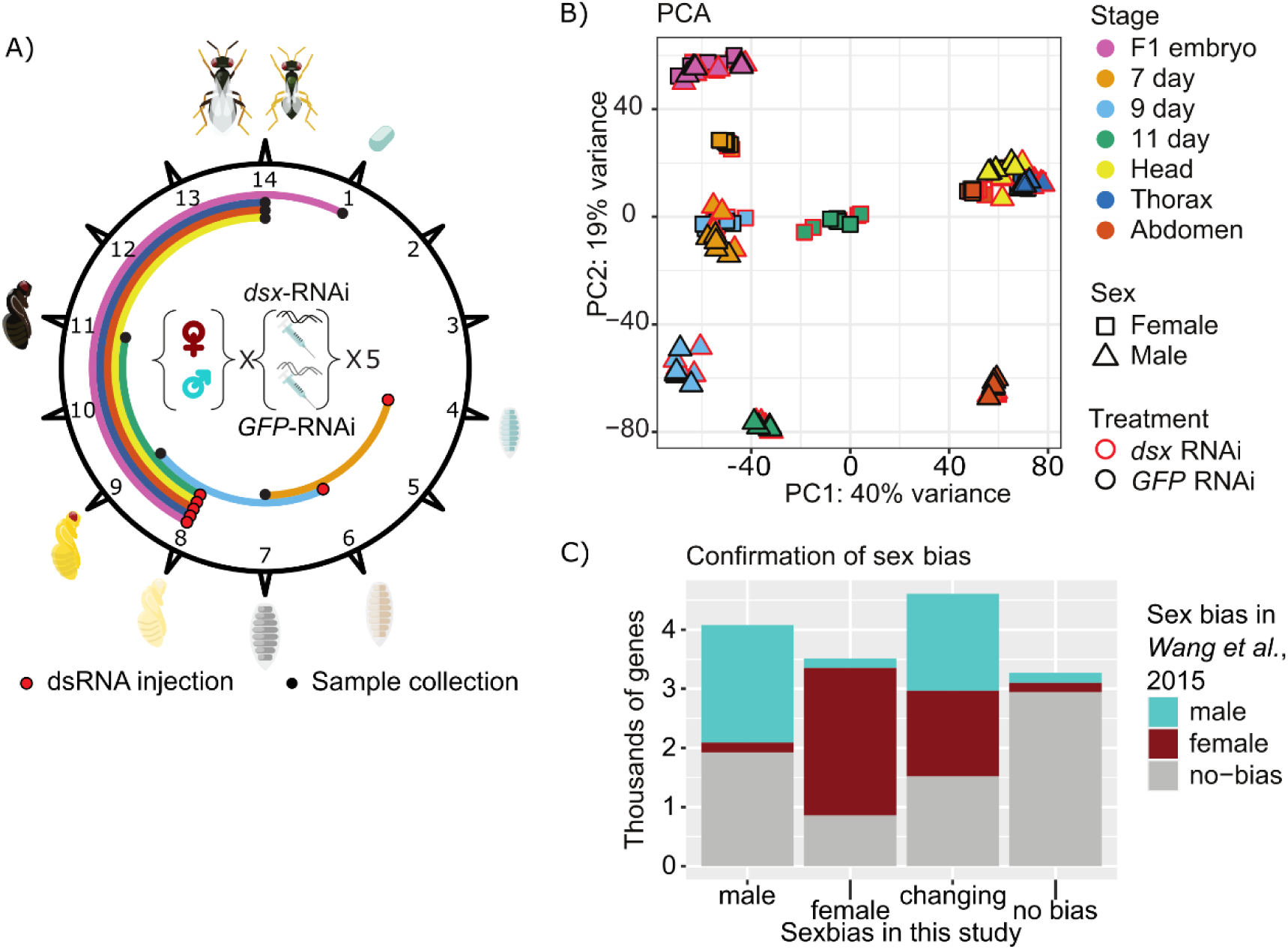
Experimental design of the transcriptomic study and data summaries. **A)** Representation of *Nasonia vitripennis* life cycle and the different times of dsRNA injection and sample collection used for the transcriptomic study. At 25°C, *N. vitripennis* develops from egg to adult in 14 days. Injection days are represented with a red dot and collection days with a black dot. The period during which the RNAi is active is represented with distinct colours for each sample type (light orange: four to seven days, light blue: six to nine days, green: eight to eleven days, yellow: eight days to adult head, dark blue: eight days to adult thorax, dark orange: eight days to adult abdomen, pink: eight days to one-day F1). Samples were collected for both males and females, injected with *dsx* or *GFP* dsRNA, in five replicates each. **B)** Principal Component Analysis for all RNA-seq samples. On the horizontal axis, PC1 represents 40% of the variance while on the vertical axis, PC2 represents 19% of the variance. Each point represents one sample. Colours depict the sample stage as presented in Fig 1A. Squares and triangles represent female and male samples, respectively. Red and black outlines depict *dsx* RNAi and *GFP* RNAi, respectively. C) Comparison of the results presented in this study with previously published data. Data from Wang *et al*. (2015) obtained in whole adults were re-analysed with the same protocol as our data. The male- and female-biased genes from this time point were compared with the sex-biased genes identified in our study. The horizontal axis represents the sex-biased genes identified in our transcriptomic analysis, while the vertical axis represents the number of genes in these categories. Changing genes are genes for which sex bias is different at various stages. Turquoise, dark-red, and grey boxes are genes found male-biased, female-biased, or not biased, respectively, in Wang et al. (2015).

#### 2.2.1 In vitro synthesis of dsRNA for RNAi

*GFP* dsRNA was generated from the vector *pOPINEneo-3C-GFP* (Addgene plasmid # 53534; https://www.addgene.org/53534/; RRID: Addgene_53534, a gift from Ray Owens). Amplification by PCR using GoTaq Flexi DNA polymerase (Promega) with GFP_RNAi_F (5’-GTGACCACCTTGACCTACG-3’) and GFP_RNAi_R (5’-TCTCGTTGGGGTCTTTGCT-3’) primers produced a 460 base-pair long amplicon covering 64% of the Emerald *GFP* CDS. dsRNA against *dsx* was generated from *Nasonia vitripennis* cDNA using the primer 5’-CCAAGAGGCAGCAAATTATG-3’ and 5’-GTTATACGCCGCATGGCTAC-3’, producing a 457 base-pair long amplicon and covering a sequence common to all male and female isoforms. Each of these PCR products was then amplified in two separate PCR to add the minimal T7 promoter sequence (5’-TAATACGACTCACTATAGGG-3’) to either end of the amplicon. The two templates were used in separate reactions to transcribe both sense and antisense RNA molecules using the MEGAscript RNAi kit (Ambion, Austin, Texas, USA) according to the manufacturer’s protocol (16 hours at 37°C). Sense and antisense RNA were mixed and annealed in equimolar amounts.

#### 2.2.2 dsRNA injections for RNAi

Larval RNAi knockdown was induced in *AsymCX* males or females at four, six, and eight days after oviposition. Larvae at four and six days were injected on PBS-1% agar plates. After injection, the four-day-old larvae were transferred back to the host (6-8 per host) and sealed with the host shell, while six-day-old larvae were kept on the PBS-agar plate. Pupae at eight days were fixed on microscopy slides with double-sided tape, injected, and incubated at 25°C until emergence. Larvae and pupae were injected in the posterior part with 4 μg/μl *dsx* or *GFP* dsRNA diluted in milliQ water mixed with red food dye in a ratio 9:1. Injections were performed using custom-made capillary needles and a Femtojet 4i (Eppendorf, 5252000021) according to the protocol by Lynch & Desplan (2006). After 72 hours, five samples containing three individuals were collected for each time point and sex and flash-frozen in liquid nitrogen. For adult parts, seven adults were collected one day after emergence and seven to eight days after pupae injection. Adults were flash-frozen in liquid nitrogen, then immediately dissected on ice to collect the head, thorax, and abdomen, and finally flash-frozen in liquid nitrogen. All samples were stored at −80 °C until RNA extraction.

For one-day-old embryos, the parental RNAi technique was used (Lynch & Desplan, 2006). Female white pupae at eight days after oviposition were injected as described above and, after emergence, crossed with non-injected AsymCX males or kept virgin. Every female was set up individually with one host, with only the anterior part accessible for oviposition. Hosts were changed every four hours. Virgin females produce only male progeny, while mated females produce 90-95% females. Batches of 100 embryos were collected 24 hours later by dissecting the host, transferred into a micro-centrifugation tube on ice without buffer, flash-frozen in liquid nitrogen, and stored at −80°C until RNA extraction.

#### 2.2.3 RNA extraction

Total RNA was extracted with Quick-RNA Tissue/Insect Kit coupled with on-column DNase treatment (ZymoResearch - R2030) according to the manufacturer’s protocol. RNA was resuspended in 10 µl DNase RNase-free water, and RNA concentration was measured on a fluorometer (Qubit 2.0, Life Technologies).

#### 2.2.4 RNA-seq library preparation

Library preparation was performed by Novogene (HK). Sequencing libraries were generated using NEBNext® Ultra TM RNA Library Prep Kit for Illumina® (NEB, USA) following the manufacturer’s recommendations, and index codes were added to attribute sequences to each sample. Briefly, mRNA was purified from total RNA using poly-T oligo-attached magnetic beads. Fragmentation was performed using divalent cations under elevated temperature in NEBNext First Strand Synthesis Reaction Buffer (5X). First-strand cDNA was synthesised using a random hexamer primer and M-MuLV Reverse Transcriptase (RNase H-). Second-strand cDNA synthesis was performed using DNA Polymerase I and RNase H. Remaining overhangs were converted into blunt ends via exonuclease/polymerase activities. After adenylation of 3’ ends of DNA fragments, the NEBNext Adaptor with hairpin loop structure was ligated to prepare for hybridisation. To select cDNA fragments of preferentially 150~200 bp in length, the library fragments were purified with AMPure XP system (Beckman Coulter, Beverly, USA). Then, 3 µl USER Enzyme (NEB, USA) was used with size-selected, adaptor-ligated cDNA at 37 °C for 15 min followed by 5 min at 95 °C before PCR. Then, PCR was performed with Phusion High-Fidelity DNA polymerase, Universal PCR primers and Index (X) Primer. Lastly, PCR products were purified (AMPure XP system), and library quality was assessed on the Agilent Bioanalyzer 2100 system. The clustering of the index-coded samples was performed on a cBot Cluster Generation System using PE Cluster Kit cBot-HS (Illumina) according to the manufacturer’s instructions. Sequencing libraries were paired-end sequenced (150 bp read-length) on an Illumina NovaSeq platform at 40M reads per library. Raw sequencing data is available at NCBI/GEO with accession number GSE260734.

#### 2.2.5 RNA-seq data analysis

Read alignment and mapping were performed by Novogene (HK). Raw data filtering was performed to remove reads with adaptor contamination, uncertain nucleotides on more than 10% of the sequence (N>10%) and reads with low-quality nucleotides (<20) on more than 50% of the sequence. Clean paired-end reads were mapped on the *Nasonia vitripennis* genome Nvit_psr_1.1 (GCA_009193385.2 Nvit_psr_1.1) using HISAT2 (v2.0.5) and default parameters. Gene counts were obtained using HTSeq (v0.6.1) and the union method on the annotation GCF_009193385.1_Nvit_psr_1. Differential expression analysis was performed in R using the software package DESeq2 (v1.28.1) (Love et al., 2014). The threshold for significant differential expression was set up at *Padj <* 0.01 when comparing time points from different sexes but the same dsRNA injection and at *Padj <* 0.05 when comparing identical sexes but different dsRNA injection. Pearson correlation analysis shows a high correlation between replicates (**Supplementary Figure 1B**). Principal Component Analysis (PCA) was performed using the built-in function from DESeq2 and displayed using ggplot2 (v3.3.5). Pearson correlations were obtained using the cor function from R (R Core Team, 2021) and displayed with ggplot2. Sankey-plot was obtained with the R package networkD3 (v0.4). Bar plots, volcano plots and violin plots on differential expression data were obtained with ggplot2 (v3.3.5). DSX regulation modes were identified using a custom R script with manual data curation in Excel.

### 2.3 DNA Affinity Purification-sequencing (DAP-seq)

We performed DAP-sequencing to identify DSX binding sites across the genome of *N. vitripennis* and tentatively indicate direct DSX targets. Briefly, in vitro produced *N. vitripennis* DSX protein is immobilised onto magnetic beads and used to pull down DNA fragments of a genomic library. Sequencing and mapping the enriched library reveal the binding sites as coverage peaks. This approach defines the DSX-binding sequence free from the influence of epigenetic information and other transcription factors. Details are provided in the next paragraphs.

#### 2.3.1 Plasmid cloning for protein production

The coding sequence for the HaloTag was PCR amplified from the plasmid pFN21K (Promega, G283A) and cloned in the TNT ICE compatible vector pF25A (Promega, L106A) via restriction by NcoI and PmeI (NEB, USA) and ligation using T4 DNA Ligase (Promega, M1804). The generated plasmid pF25A-HaloTag was transformed in TOP10 competent *Escherichia coli* (Invitrogen; C404003). The coding sequence for *Nasonia vitripennis* DSX^F^ (Dbxref: GeneID:100302336, Genbank:NP_001155990.1) was obtained as a long oligo from Eurofins genomics and subsequently cloned downstream the Halotag in the pF25A-HaloTag via PCR amplification, restriction with AsisI and PmeI (NEB, USA), and ligation using T4 DNA Ligase (Promega, M1804). Similarly, the luciferase coding sequence was PCR amplified from the control vector Luciferase ICE T7 Control DNA (Promega, L105B) and cloned downstream the HaloTag in the plasmid pf25A-HaloTag. The pf25A-HaloTag-NvDSXF and the pf25A-HaloTag-Luciferase vectors were transformed in TOP10 competent *E. coli* (Invitrogen; C404003) for propagation. The vectors used are available upon request.

#### 2.3.2 DNA library preparation

DNA libraries were produced by shearing 1 μg of purified male and female genomic DNA using a Covaris ultrasonicator (duty cycle: 10%, intensity: 5%, cycles/burst: 200, time: 60s, number of cycles: 3) to obtain fragments 200bp to 400bp long. Sodium acetate purification followed. Purified fragments were repaired and A-tailed using the NEBNext® Ultra™ II DNA Library Prep Kit for Illumina® (NEB, E7645S). Another round of sodium acetate purification followed. Universal, Y-shaped adaptors were ligated to the DNA fragments using T4 DNA Ligase (Promega, cat. no. M1804). Another round of sodium acetate purification followed. These libraries were subjected to protein-mediated enrichment, as described below.

#### 2.3.3 Protein production and library enrichment

DSX^F^ and Luciferase proteins were produced *in vitro* using TnT® T7 Insect Cell Extract Protein Expression System (Promega, L1101). Production of protein was assessed by dotblot with the use of anti-HaloTag antibodies (Promega, G9211) and AP-conjugated secondary antibodies (Promega, S3721). After four hours of incubation, proteins were immobilised on magnetic beads (Promega, G7281) by continuous mixing for two hours. Three washes with PBS - 0.005% NP40 were used to remove non-tagged proteins. A library of 75 ng of DNA was added to each tube, and a two-hour incubation on continuous rotation followed. After incubation, four washes using PBS - 0.005% NP40 were done to remove nonspecific-interacting DNA. The DNA was eluted in EB buffer (Qiagen, Cat. No. / ID: 19086) by denaturing the proteins at 98°C for 10 minutes. Recovered, enriched DNA libraries were subjected to 20 PCR-amplification cycles using uniquely barcoded primers. Amplification products were subjected to gel electrophoresis, and a gel slice corresponding to fragments of size between 200 and 400 bp was excised; DNA purification via gel extraction kit followed (Qiagen; QIAquick Gel Extraction Kit cat no. 28704). Each library was sequenced by Novogene (HK) on a Novaseq Illumina platform, paired-end 150bp with 20M reads as target sequencing depth.

#### 2.3.4 DAP-seq data analysis

Sequences were aligned on Nvit_psr_1.1 genome (GCA_009193385.2 Nvit_psr_1.1) using BOWTIE2 (v2.3.5.1) (Langmead & Salzberg, 2012). Coverage peaks were identified using the function callpeak - g 2.96e8 -q 0.01 in MACS2 (v2.2.5) (Zhang et al., 2008), pooling the four DSX-enriched libraries aligments and comparing them with four Luciferase-enriched libraries alignment. Sequences in MACS2-defined peaks were retrieved to find de novo binding sites using the function meme-chip -ccut 0 -dna – o from MEME (v5.1.0) (Machanick & Bailey, 2011). The raw sequencing data and the peaks identified with MACS2 can be downloaded from GEO/NCBI from the accession number GSE284068. The obtained minimum consensus sequence was used to scan the genome for binding sites. In particular, we retrieved gene regions defined as the sequence of each gene plus five kilobases upstream and one kilobase downstream the sequence of each gene annotated gene in the Nvit_psr_1.1 genome, and we counted the number of DSX binding sites per gene region.

## 3 Results

### 3.1 Overview of the transcriptomic approach

We compared gene expression of male and female *N. vitripennis* upon RNAi-mediated knockdown of *dsx* or, as a negative control, the exogenous gene *Green Fluorescent Protein* (*GFP*). For each treatment, we collected samples at one, seven, nine, eleven and fourteen days of development, representing respectively embryos, late larvae, early pupae, late pupae and adult life stage. From adult samples, we separately collected heads, thoraces, and abdomens. This experimental design is summarised in **Figure 1A**. We verified that all *dsx-*silenced adult samples had phenotypic changes, as shown by Wang et al. (2022a), and the *GFP-*silenced adults did not. Each sample consisted of five replicates and was paired-end sequenced with Illumina technology at 40M reads per library.

A similar number of genes is expressed in each sample (**Supplementary Figure 1A**); therefore, throughout the analysis and in this publication we report the number of sex-biased genes in relation to the total number of genes in the current annotation (GCA_009193385.2 Nvit_psr_1.1).

Pearson correlation analysis shows a high positive correlation between replicates (**Figure 1B**). Principal Component Analysis (PCA) corroborates the clustering of replicates and shows that the developmental stage or adult body part explains most gene expression variation. Moreover, samples cluster by sex, with a progressive increase in differences during development (**Figure 1B**). Transcriptomes are most similar in the F1 embryos and become less similar throughout development, with the eleven-day-old pupae showing the highest variation between the sexes. In adult head and thorax, male and female samples cluster together while abdomen samples show clear differential clustering between sexes. Surprisingly, *dsx* RNAi does not cause strong shifts in gene expression profiles.

### 3.2 Identification and confirmation of sex-biased gene expression across development

We first analysed the *GFP* (control) samples to determine the extent and distribution of sex-biased gene expression and compared this with previously published RNA-seq data of male and female adults (Data ref: Wang et al., 2015). We compared the sexes for each developmental time point and identified a cumulative total of 78.9% (12,199/15,471; *Padj <* 0.01) sex-biased genes. Then, we re-analysed the *N. vitripennis* data of Wang et al., (2015) with our analysis protocol and found 57.0% (8,797/15,471) sex-biased genes in whole adult bodies, with 4,347 male-biased and 4,450 female-biased genes (*Padj <* 0.05) in their dataset. A comparison of these datasets shows a high correlation between the gene expression profiles of males and females, corroborating the previous results by Wang et al. (2015) that highlighted the vast extent of differential gene expression between sexes (**Figure 1C**). Due to the higher number of sample types in our dataset, we identified 1,924 new male-biased genes and 859 new female-biased genes that were previously not identified as sex-biased.

### 3.3 Sex-biased gene expression is dynamic and increases as development progresses

Sex-biased gene expression increases sharply over developmental time, starting at 393 (2.5% of all annotated genes) sex-biased genes in embryos, increasing to 5,076 (32.8% of all annotated genes) in 11-day-old pupae (three days before emergence), and reaching a total of 11,260 genes across all adult body parts (72.8% of all genes) (**Figure 2A**). We report this information in detail as MA plots in **Supplementary Figure 2,** and as machine readable data in **Supplementary Table 1**. The number of sex-biased genes is higher in female than male embryos (**Figure 2A)**. In pupal stages at 7 and 9 days, the number of sex-biased genes is higher in males than females (**Figure 2A**), while it is similar in 11-day-old pupae (**Figure 2A**).

**Figure 2:**
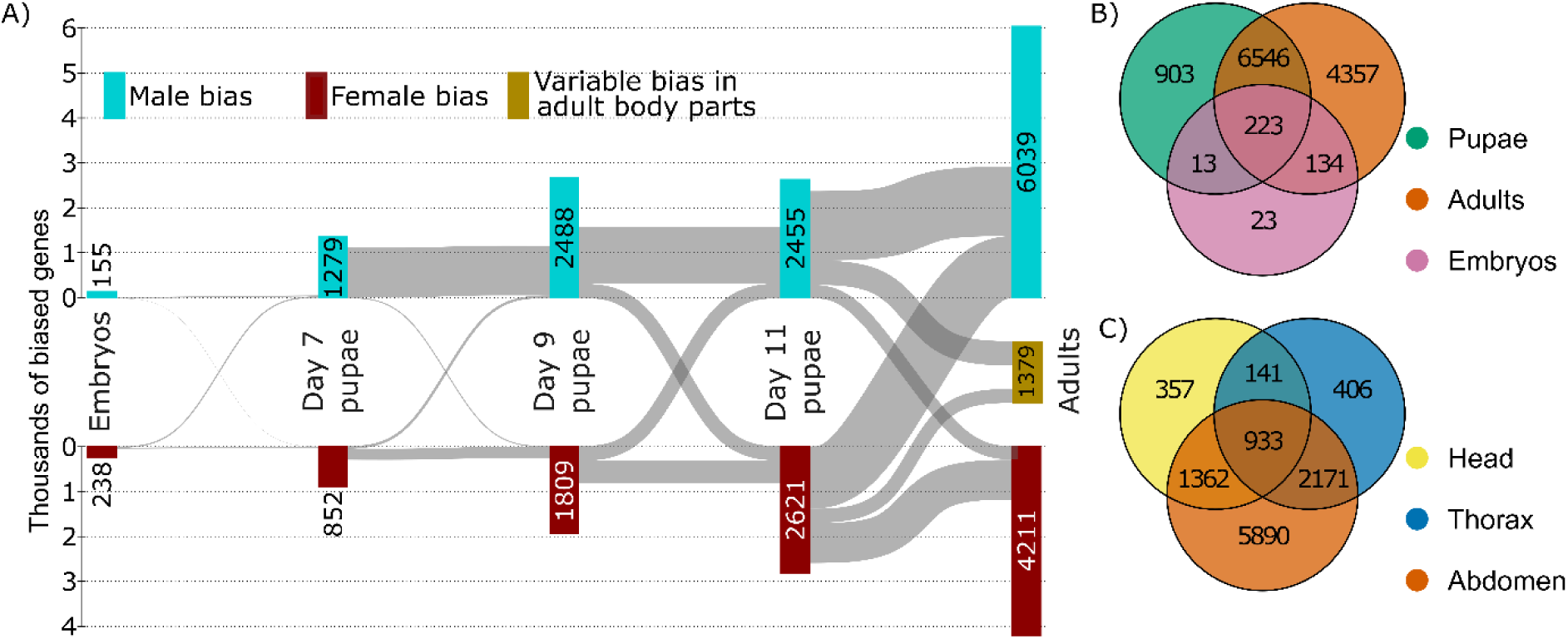
Sex-biased gene expression progression during *Nasonia vitripennis* development. **A)** Sankey diagram showing the number of male (turquoise edge box) and female (red edge box) biased genes at each stage in development and its progression to the next developmental stage represented by paths connecting boxes. The adult stage shows the combined values of all three adult body parts. Genes with different sex biases in different adult body parts were categorised as variable (black edge box). Box and path sizes are proportional to the number of genes they represent. **B)** Venn diagram presenting the overlap between sex-biased genes in embryo (pink), pupal (green), and adult (orange) samples. For pupal samples, the combined sex-biased genes at 7, 9 and 11 days after oviposition were used. For adult samples, the combined sex-biased genes in all three body parts were used. **C)** Venn diagram presenting the overlap between sex-biased genes in adult head (yellow), thorax (blue), and abdomen (orange) samples.

Interestingly, 88% (6,546/7,685) of sex-biased genes in the pupal stages are also sex-biased in adults, and 39% of all adult sex-biased genes (4,357/11,260) are exclusive for adults (**Figure 2B**). Only 223 genes have a sex-biased expression in embryos and at least one pupal and adult sample (**Figure 2B**). In the separate adult tissues, 80% of the sex-biased genes in the head (2,295/2,853) and 85% (3,104/3,651) of the sex-biased genes in the thorax are also sex-biased in the abdomen, while 57% of all abdominal sex-biased genes (5,890/10,356) are exclusive for the abdomen: as a result, the highest number of sex-biased genes (10,356) is found in the adult abdomen (**Figure 2C**). Once genes become sex-biased, they mostly remain sex-biased in the next developmental stage (**Figure 2A**) and retain their male or female bias. Nonetheless, some reversal of sex-biased expression is also observed: in females, transitioning from 11-day-old pupae to adults causes 1102 genes to become male-biased, while 1519 genes remain female-biased (**Figure 2A, Supplementary Table 1**).

In conclusion, sex-biased expression during *N. vitripennis* development is highly dynamic, and the number of sex-biased genes increases during development. The extent of sex-biased expression also depends on the tissue analysed, with abdomens showing the highest divergence.

### 3.4 DSX controls the expression of sex-biased genes

After establishing differentially expressed genes (DEGs) between sexes, we sought to determine to what extent DSX is responsible for it. To this end, we compared transcriptomes of *dsx* knockdown individuals with those of control individuals per time point and sex (**Figure 3A**, **Supplementary Table 1**). In total, in all groups and both sexes, 64.6% of all sex-biased genes (7,880/12,199) appear to be under DSX transcriptional control as these were either significantly up- or down-regulated after *dsx* knockdown: this is more than expected by chance alone (χ² (1) = 2540.7, p < 2.2e-16). Second, we found that *dsx* knockdown mainly affects gene expression in embryos and adults and has minimal effects on gene expression in larval and pupal stages (**Figure 3A**). Third, *dsx* knockdown has a stronger effect on gene expression in males than in females, except for the embryonic stage where DSX regulates 15.2% of all genes in females and 1.9% of all genes in males. Finally, we do not observe a tendency for predominant up- or down-regulation after *dsx* knockdown, but rather a general deregulation of sex-biased gene expression. This suggests that DSX acts both as an activator and a repressor of transcription.

**Figure 3:**
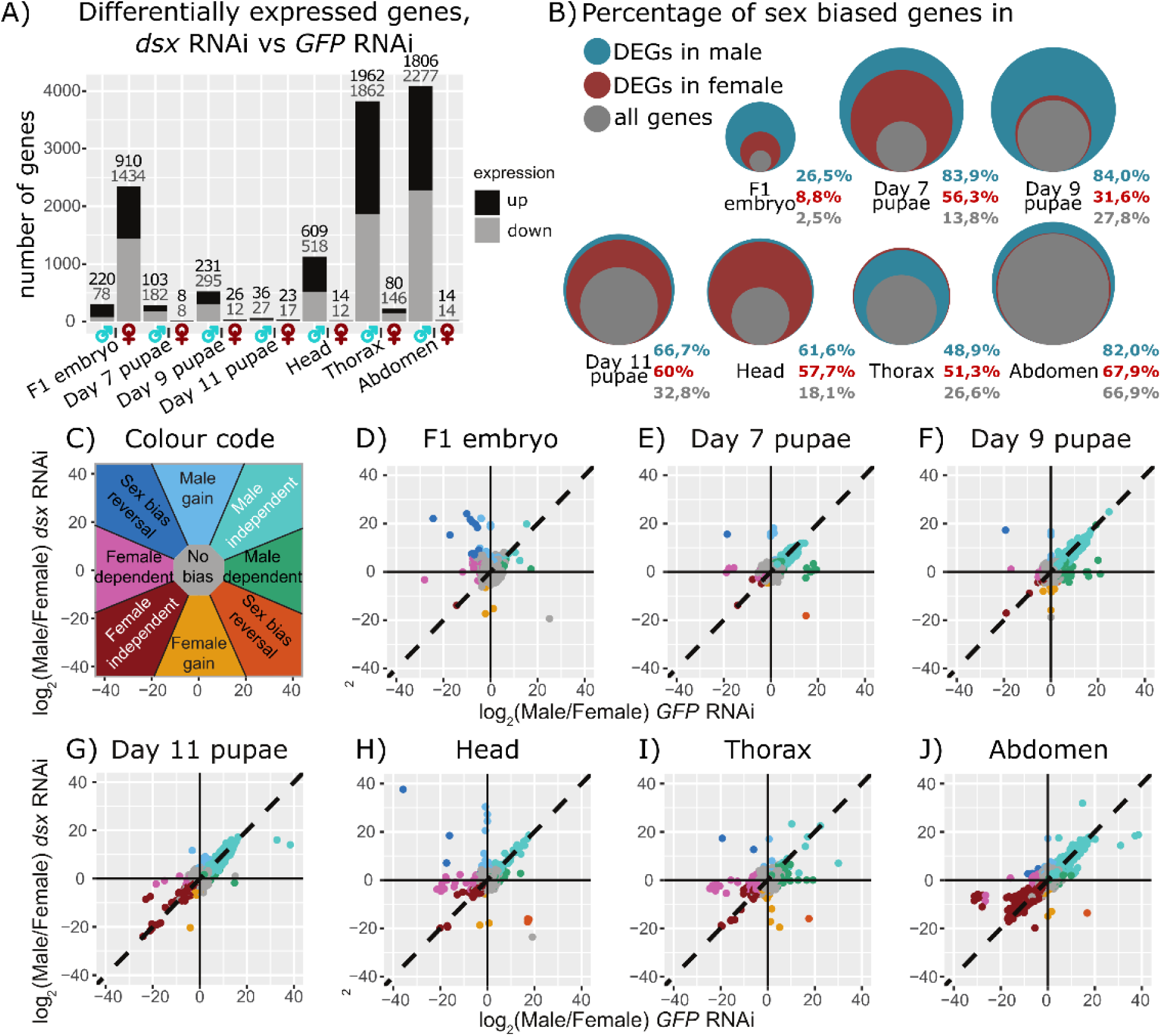
*dsx* controls sex-biased gene expression using different modes of regulation. **A)** Number of differentially expressed genes (DEGs) between *dsx*-RNAi and *GFP*-RNAi samples for all stages and both sexes. Black: log2FC(*dsx*-RNAi/*GFP*-RNAi)>0 and *Padj < 0*05; grey: log2FC(*dsx*-RNAi/*GFP*-RNAi)<0 and *Padj < 0*05. Above each stacked bar we report the number of genes up- or down-regulated upon *dsx* RNAi. **B)** Percentage of sex-biased genes (as identified in Figure 2A) among the DEGs identified in Figure 3A. Circle areas are proportional to the percentage of sex-biased genes for each gene subset. Grey: Percentage of sex-biased genes among all genes in the annotation. Turquoise: percentage of sex-biased genes within the male DEGs. Red: percentage of sex-biased genes within the female DEGs. **C)** Model for the dot plot interpretation of the relative fold-change comparison between *dsx* RNAi and *GFP* RNAi gene expression groups. Genes with a strong male- or female-biased expression that lose their sex bias by *dsx* knockdown fall in the male-dependent or female-dependent sections, respectively and are dependent on *dsx* for their sex-biased gene expression. Those genes that change in their sex-biased expression as a result of *dsx* knockdown fall in the sex-bias reversal sections, and those that maintain their sex bias are female or male-independent and are seemingly independent of *dsx* regulation. Non-biased genes that gain a sex bias fall into the male- or female-gain sections, and non-biased genes that are not responding to *dsx* knockdown fall in the middle. Colour code corresponds to dot colours in D-J. **D-J)** Dot plots showing for each stage the fold change (log2-transformed) between *dsx* knockdown male and female samples as a function of the fold change (log2-transformed) between *GFP-*dsRNA injected male and female samples (control). For both conditions, samples are considered male-biased when log2FC(male/female)>0 and *Padj <* 0.01, and female-biased when log2FC(male/female)<0 and *Padj <* 0.01. The colour code is as follows: turquoise: male bias in both control and treatment; green: male bias in control and no bias in treatment; orange: male bias in control and female bias in treatment; yellow: no bias in control and female bias in treatment; red: female bias in both control and treatment; pink: female bias in control and no bias in treatment; dark blue: female-bias in control and male bias in treatment; light blue: no bias in control and male bias in treatment, grey: no bias in both control and treatment.

We then investigated whether DEGs after *dsx* knockdown are enriched in the sex-biased gene sets: an enrichment would confirm that DSX controls sex-biased gene expression. We found that at each time point, there is an enrichment of sex-biased genes in the DEG subsets for both males and females (**Figure 3B**). By plotting the relative fold-change of gene expression in *dsx* knockdown individuals against gene expression in the control condition for each group we can infer the effect of DSX on sex-biased expression, i.e. if it is increasing or decreasing bias and towards the male or female direction (**Figure 3C**). The main effect of *dsx* knockdown is a decrease in both male and female sex bias (**Figure 3D-J**). Across all time points, 78 genes show reverted sex-biased expression after *dsx* knockdown, with female-biased genes becoming male-biased and vice versa. This phenomenon is most pronounced in embryos, where 23 female-biased genes become male-biased, while the reverse effect never occurs (**Figure 3D**). Moreover, 579 genes that were previously not sex-biased, gained sex-biased expression upon *dsx* knockdown in at least one time point, especially in embryos, 11-day-old pupae, head, and abdomen samples. Altogether, the global changes in gene expression during development after *dsx* knockdown show that DSX acts as a master regulator of sex-biased expression through a complex and dynamic process for more than 50% of all *N. vitripennis* genes.

### 3.5 DSX acts mainly as an activator or repressor in males

We confirmed that DSX controls sex-biased gene expression, yet its mode of action on gene expression remains to be explored. Many possibilities of gene expression control can explain the observed sex-biased expression and are summarised in **Supplementary Figure 3A** and 3B: for example, DSX could be an activator, a repressor, or have opposite effects in males and females. To explore the DSX mode of gene-expression control, the sex-biased gene expression was compared with the effect of *dsx* knockdown in both males and females at every time point separately; this is visualised in **EV Figure 1A**. In this way, the DSX mode of gene regulation could be deduced unequivocally. As an example, this analysis for male and female thorax samples is visualised in **EV Figure 1B**, and the other analyses are presented in **Supplementary Figure 4.** The data from every time point were then merged to obtain a global table explaining the sex-bias expression for every gene and the DSX mode of action (**Figure 4A**, **Supplementary Table 2**).

**Figure 4:**
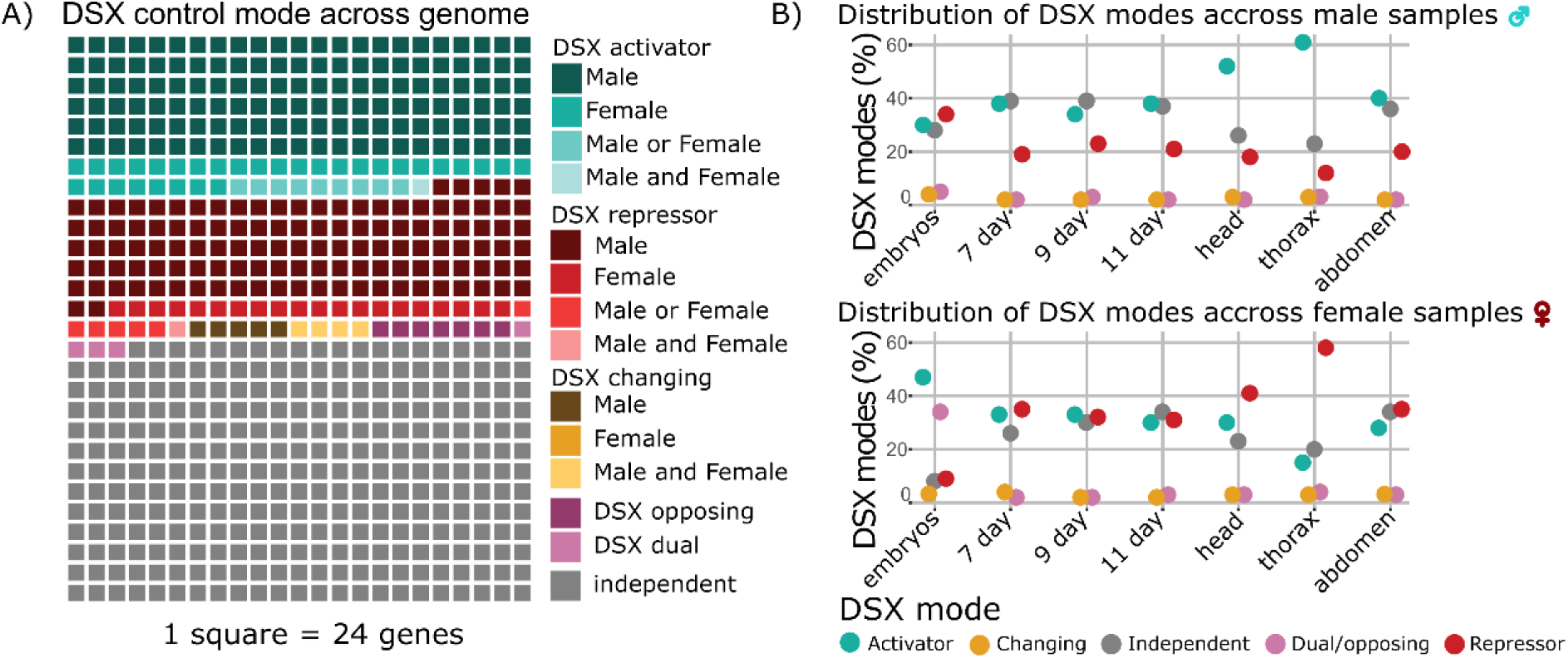
DSX modes of gene expression regulation. **A)** Waffle tree showing DSX gene control mode distribution across the whole genome after analysis of all seven sample types. Each square represents 24 genes. The number of genes for each mode was rounded to the closest multiple of 24. **B)** Dot plot showing DSX gene expression control modes for each sample type, regardless of the sex bias. DSX modes are ranked per abundance. Numbers represent the percentage of genes controlled by each DSX mode for each sample type. The colour code is as follows: red: repressor, aquamarine: activator, ochre: changing, pink: dual/opposing, dark grey: independent of DSX control.

We found that 54.1% of all genes in our annotation have a DSX mode of action, while the remainder seemed independent of DSX regulation. Furthermore, all modes of control are present, yet DSX acts mainly in males by activating or repressing genes **(Figure 4C, Supplementary Table 2):** there are five times more genes controlled by DSX in males than in females. Cases of dual, opposing effects (activation in one sex and repression in the other) are rare and concern only 270 genes.

Comparing the distribution of DSX modes of gene regulation across time points and sex (**Figure 4B**) shows that genes activated by DSX are overrepresented in the male-biased gene set from 11 days onwards, while genes repressed by DSX are found mainly in the female-biased genes. There is also a remarkable difference in DSX mode repartition for female-biased genes in embryos, as it is the only stage where female-biased expression is explained by an activator or a dual function of DSX.

We now have a clear overview of which genes are sex-biased and how they can be controlled by DSX. However, it is not clear whether these genes are direct or indirect targets of DSX regulation. We explored this question by means of a DNA-protein interaction technique to provide information on DSX binding on *N. vitripennis* genome.

### 3.6 DSX binds a conserved DNA motif

To identify the direct targets of DSX, we set out to identify which genes contain a DSX binding site in their regulatory region. However, we first needed to determine the DSX binding sequence. To this end, we performed in vitro DNA-affinity purification coupled with sequencing (DAP-seq) (Bartlett et al., 2017).

Combining four DSX-enriched libraries, we identified 2307 peaks. *De novo* motif analysis revealed 903 occurrences of a putative binding site. The identified consensus sequence, [AT]TACA[TA]TGT[TA][TG][CT], is similar to the *D. melanogaster* DSX binding sequence reported by Luo et al., (2011) (**Figure 5A**). The probability matrix for each nucleotide is provided in **Supplementary Table 3**.

**Figure 5:**
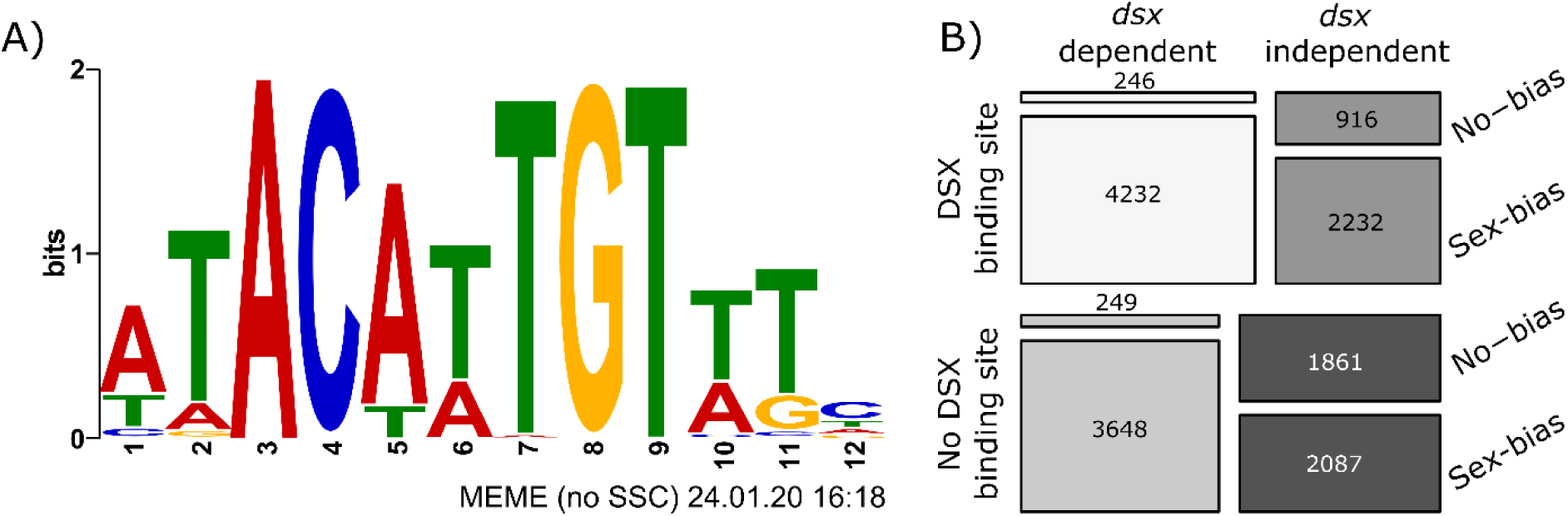
Characterisation of DSX binding sites and control on target genes. **A)** WebLogo motif derived from the alignment of sequencing library enriched with DSX bait in DAP-seq experiment. On the X-axis is the position of the nucleotide in the motif, and on the Y-axis is the information content in bits. **B)** Mosaic plot showing the overlap between sex-biased and non-sex-biased gene expression, *dsx*-dependent or independent control of expression, and presence or absence of DSX binding sites for all genes in the annotation. In total, 4232 genes are sex-biased, controlled by DSX, and have at least one DSX-binding site. The observed values for all categories (tiles) diverge significantly from what is expected based on independent observations (standardised residuals bigger than 4 or smaller than −4).

Luo et al. (2011) proposed that the optimal DSX-binding sequence is enriched in the *Drosophila melanogaster* genome by virtue of DSX pleiotropic regulatory function. We established that DSX regulates thousands of genes in *N. vitripennis;* therefore, we expected a similar enrichment in DSX-binding sites in the *N. vitripennis* genome. Indeed, all ten sequences with the most hits for DSX binding are represented in the genome more than expected by chance (**Supplementary Table 4**). Notably, sequences that diverge from the optimal DSX-binding sequence are also represented in the DAP-seq dataset, suggesting that DSX could target these sequences in vivo.

### 3.7 Candidate genes under direct DSX control

The DSX**-**binding-site motif with the relaxed identity (NN[TA]ACA[AT]TGT[AT]NN) occurs in the *N. vitripennis* genome 12,786 times. We identified 7,687 genes (49.6% of all genes) with at least one such DSX binding site in their regulatory region, defined as the window between 5 kb before the transcription start site and 1 kb after the transcription end site.

By combining sex-biased expression, DSX mode of control, and the presence of a DSX-binding-site motif, we identified 4,232 potential direct DSX target genes (27.3% of all genes) (**Figure 5B**). We found that genes containing the DSX-binding-site motif are enriched in sex-biased genes compared to non-target genes (χ² (1, N = 15,274) = 233.0525, p < 2.2e-16) (**Supplementary Table 1**) (**Figure 5B**). Moreover, genes with one binding site are enriched in male (749 observed over 693 expected) and female bias (672 observed over 598 expected), whilst genes with more than one binding site are enriched in genes with changing sex bias (524 observed over 330 expected) (χ² (6, N = 15,274) = 230.7, p < 2.2e-16) (**Supplementary Figure 5A**). This suggests that DSX can bind to different enhancers to achieve sex-, time-, and tissue-specific gene regulation. We found that genes containing the DSX binding site motif are enriched in genes controlled by DSX (χ² (1, N = 15,274) = 127.1, p < 2.2e-16). When dividing genes into categories with either zero, one or more than one DSX-binding site, there is a significant enrichment of “activated gene” in the subset of genes with more than one binding site (1283 observed over 993 expected) and enrichment of “independent genes” in genes without DSX binding sites (3754 observed over 3412 expected) (χ² (8, N = 15,274) = 222.6, p < 2.2e-16) (**Supplementary Figure 5B**).

In *D. melanogaster*, 3,717 genes (23.5% of all genes) were described as potential direct DSX targets (Clough et al., 2014), of which 1,533 genes have a direct homolog in *N. vitripennis.* Among these genes, 957 (63.1%) genes have a DSX-binding site in *N. vitripennis*, which is more than expected by chance (χ² (1, N = 3,066) = 50.3498, p < 0.00001) (**Supplementary Table 1**). These shared genes are enriched in sex-biased genes, especially within the category-changing sex-bias, and DSX-controlled genes, especially with DSX as an activator (**Supplementary Figure 5C, D**). Within these genes, well-known DSX targets are present, such as several *yellow* genes (*yellow-y*, *yellow-f*, *LOC100116775*), *bric-à-brac* genes (*LOC100123373*, *LOC100124164*), *vitellogenin* (*LOC100114046*), *Abdominal-B* (*LOC100119890*), *fruitless*, and *dsx* itself.

### 3.8 DSX control of noteworthy genes

In addition to describing the general mode of gene regulation, our dataset provides a map to explore *dsx* effects on each gene. We selected a few sex-biased, well-characterised genes to benchmark our dataset and analysis.

Genes acting upstream of *dsx* in the sex-determination cascade are well-characterised in *N. vitripennis*. The two duplicated genes *womA* and *womB*, are not expressed in any sample (**Supplementary Table 1**), as we would expect based on its defined temporal expression pattern limited to early embryos (Zou et al., 2020). The gene *tra* is one of the three genes that are female-biased at all time points and in all tissues (adj. p value < 0.01; between 2.93 and 8.51-fold higher in females than males) (**EW Figure 2A**). Moreover, its expression is not affected by *dsx* knockdown, except for a mild increase in *dsx* knockdown female embryos (adj. p value < 0.01; 1.32-fold increase). The gene *Tra2* (LOC100116671) is expressed at various levels across development, is female-biased in the abdomen and the head (adj. p value < 0.01; 1.61 and 1.21-fold, respectively), and is downregulated by *dsx* knockdown in the male thorax and abdomen (adj. p value < 0.01; 0.80 and 0.85-fold, respectively) (**EW Figure 2B**).

Another well-characterised sex-biased gene is *vitellogenin*, which codes for a family of proteins that nurture developing eggs and embryos (Raikhel & Dhadialla, 1992). It is expressed in female fat bodies and has been shown to be under DSX control in a wide range of insect orders (Chikami et al., 2022; Shukla & Palli, 2012; Suzuki et al., 2003; Wexler et al., 2019). *N. vitripennis* has two copies of *vitellogenin* (LOC100122102 and LOC100114046) located only 4 kb apart: both have female-biased expression in 11-day-old pupae and all adult tissues (**EW Figure 2C, D**). Upon *dsx* knockdown, their expression strongly increases in males and remains constant in females. In particular, the expression of LOC100122102 is between 300 and 14,000 times higher in females compared to males (adj. p value <0.01), and *dsx* knockdown causes its expression to increase between 3,000 and 9,000 times in males (adj. p value <0.01). LOC100114046 is expressed between 12 and 55-fold in females compared to males (adj. p value <0.01), and *dsx* knockdown causes its expression to increase between 16 and 26-fold in males (adj. p value <0.01).). The remarkable similarity in the deregulation of the two *vitellogenin* genes upon *dsx* knockdown suggests that a single enhancer (or the same, duplicated enhancer) could be involved in the regulation of both genes. Both LOC100122102 and LOC100114046, have DSX binding sites in the 5 kilobase region upstream their respective transcription start site (**EW Figure 2F)**

Vitellogenins localise into the developing eggs via receptor-mediated endocytosis (Raikhel & Dhadialla, 1992). Therefore, vitellogenin receptor genes are often investigated as possible DSX targets. As expected, the vitellogenin receptor gene LOC100119119 is strongly female-biased in adult abdomens (adj. p value < 0.01; 7.84-fold) (**EW Figure 2E**). Upon *dsx* knockdown, only a slight decrease in the male abdomen and female embryo is detected (adj. p value < 0.01; 0.66 and 0.68-fold, respectively). In the proximity of LOC100119119, there are two putative DSX binding sites: CCTACAATGTATT and TTGACATTGTGAC are located at 2824-2811 bp and 3806-3793 bp before the start of the transcription site, respectively (**EW Figure 2F**). The effect of *dsx* knockdown on LOC100119119 is mild because the formation of male and female reproductive organs is not affected by *dsx* deregulation at the pupal stage but rather in the early larval stages, as shown by Wang et al. (2022a).

## 4 Discussion

Combining gene expression analysis to define sex-biased genes and genes affected by *dsx* knockdown with the DSX binding-site sequence analysis provided a model for DSX control on *N. vitripennis* sex-specific transcription. We show that 78.9% of all *N. vitripennis* genes exhibit sex-biased expression at some point during development. This is almost 40% more than could be identified from the previously published data by Wang et al. (2015). Moreover, 27.3% of all genes are likely under direct DSX control and 37.7% are under indirect DSX control. In males, DSX regulates five times as many genes as in females, and the majority of these genes are activated by DSX. In contrast, the DSX-regulated genes in females are mainly repressed. Our dataset provides a solid framework for research on sexual differentiation in Hymenoptera and DSX regulatory evolution across insect orders.

We find that the number of sex-biased genes increases over time, but the number of expressed genes does not significantly change. From this, we infer that sex-biased expression results from genes acquiring sex-specific bias during development rather than specific subsets of genes being expressed solely during late developmental stages. Moreover, most sex-biased genes retain their sex-specific bias across developmental stages, especially in males. However, many female-biased genes become male-biased at the transition from pupal to adult stages. This category of genes is of particular interest to explore interactions between DSX and other transcription factors that could control the spatiotemporal nature of sex-biased gene expression.

Knockdown of *dsx* strongly affects males during larval and adult stages, while it has a more subtle effect on female at these developmental stages; this aligns with previous observations relative to male phenotypic feminisation (Wang et al., 2022a). In contrast, *dsx* knockdown in one-day-old embryos affects the female transcriptome more strongly, suggesting a difference in the sex-bias regulation mode in early development. Knockdown of *dsx* also has a more significant impact on adults than on pupae, possibly due to the longer interval between dsRNA injection and sample collection, allowing for potential indirect effects on gene expression or because adults naturally exhibit a more significant number of sex-biased genes.

We used RNAi to transiently knock down *dsx* and collected samples 72 hours after treatment. In this way, we likely captured only a limited picture of the genes regulated by DSX. An example is the lack of *dsx* knockdown effects on the *vitellogenin receptor* gene expression, despite the known role of DSX in testes and ovary development (Wang et al., 2022a), and highlights a potential limitation of our study in detecting genes indirectly regulated by DSX.

The identified *N. vitripennis* DSX binding sequence shares a conserved core with *D. melanogaster* DSX and the mammal DSX homolog DMRT1 (Luo et al., 2011; Murphy et al., 2010). This strongly suggests that the DSX-binding sequence is conserved in insects. Nonetheless, the peripheral nucleotides diverge considerably from the ones identified in *D. melanogaster*. However, DSX is known to bind and regulate enhancers with sequences that diverge considerably from the best affinity DSX binding sequence. In *D. melanogaster* yolk protein regulation, DSX recognises two binding sites that share 11/13 and 7/13 nucleotides with its highest affinity sequence (Burtis et al., 1991; Luo et al., 2011). These deviations are necessary to integrate sexual, spatial, and temporal information into the regulation of yolk protein expression: the DSX binding site overlaps with the binding site for other transcription factors, facilitating enhanced expression in fat bodies and repressed expression in ovaries (An & Wensink, 1995). Similar “sequence compromises” are also expected to apply to *N. vitripennis’* regulome.

By merging data on sex-biased gene expression, the impact of *dsx* knockdown in males and females, and the identification of the DSX-binding sequence, we determined the regulatory role of DSX for each gene. In particular, we defined whether DSX is active in males, females, or both sexes and whether DSX functions as an activator, repressor, dual transcription factor, or exhibits dynamic activity changes over development and tissue. The initial mode of DSX regulation identified in *D. melanogaster* with an antagonistic effect in males and females is observed in *N. vitripennis* as well but appears to be rare. The most common modes in which DSX exerts control are, in order of decreasing frequency, as an activator in males, as a repressor in males, as a repressor in females and as an activator in females.

The male-centric role of DSX seems conserved in Hymenoptera. In *Athalia rosae*, *dsx* depletion throughout male development causes complete feminisation but does not affect female development (Mine et al., 2021). Knocking out *dsx* in *Apis mellifera* females results only in ovary deformities, suggesting that DSX has a limited feminisation effect in this species (Roth et al., 2019). The effects of *dsx* knockout on male development have not been investigated but are expected to be more severe and pervasive. This male-centric DSX control of sex differentiation is confirmed to be a plesiomorphism, as it is shared in basal insect orders such as Zygentoma (Chikami et al., 2022) and Blattodea (Wexler et al., 2019), where the role of DSX in females appears to be limited to a cryptic feminisation function, involving only a few genes, mainly *vitellogenins*. In Diptera, Coleoptera and Lepidoptera, collectively known as Aparaglossata, DSX has an essential role in feminisation. Chikami et al. (2022) propose that the differences in female transcriptome regulation between the Aparaglossata and the other insects arise from the evolution of a disordered C-terminal domain in the DSX female isoform. Nevertheless, the effects of DSX on the female transcriptome of most insect species are probably not fully recognised. Most studies focus on gene expression in adults, yet we show that 90% (1122/1239) of the genes for which DSX acts as a repressor or activator in females are regulated by DSX at the embryonic stage.

Altogether, our study provides a second model for DSX’s role in regulating sexual dimorphism, in addition to the one developed for *Drosophila* (Clough et al., 2014; Hildreth, 1965). By identifying similarities in the two models, such as the common DSX-DNA-binding sequence and the double DSX activity as an activator and a repressor, we provide a generalised model applicable to most insects. By identifying peculiarities of our model, such as the male-centric role of DSX regulatory activity, we provide a model that could explain sexual dimorphism regulation in insects where DSX has a predominant role in masculinisation.

## Supporting information

TableS1_data_summary

Supplementary_collection

## Acknowledgements

This work was by the Dutch Research Council (NWO).

We would like to thank Marcel Dicke for precious comments on a previous version of this paper.

## Author contribution

F. Guerra: Conceptualization; Validation; Investigation; Data curation; Visualization; Writing—original draft; Writing—review and editing.

J. Rougeot: Conceptualization; Validation; Investigation; Data curation; Visualization; Writing—original draft; Writing—review and editing.

E.C. Verhulst: Supervision; Funding acquisition; Project administration; Writing—review and editing

