## Supplementary_collection for "A transcriptional control model for *doublesex*-dependent sex differentiation in *Nasonia* wasps"

### Expanded view

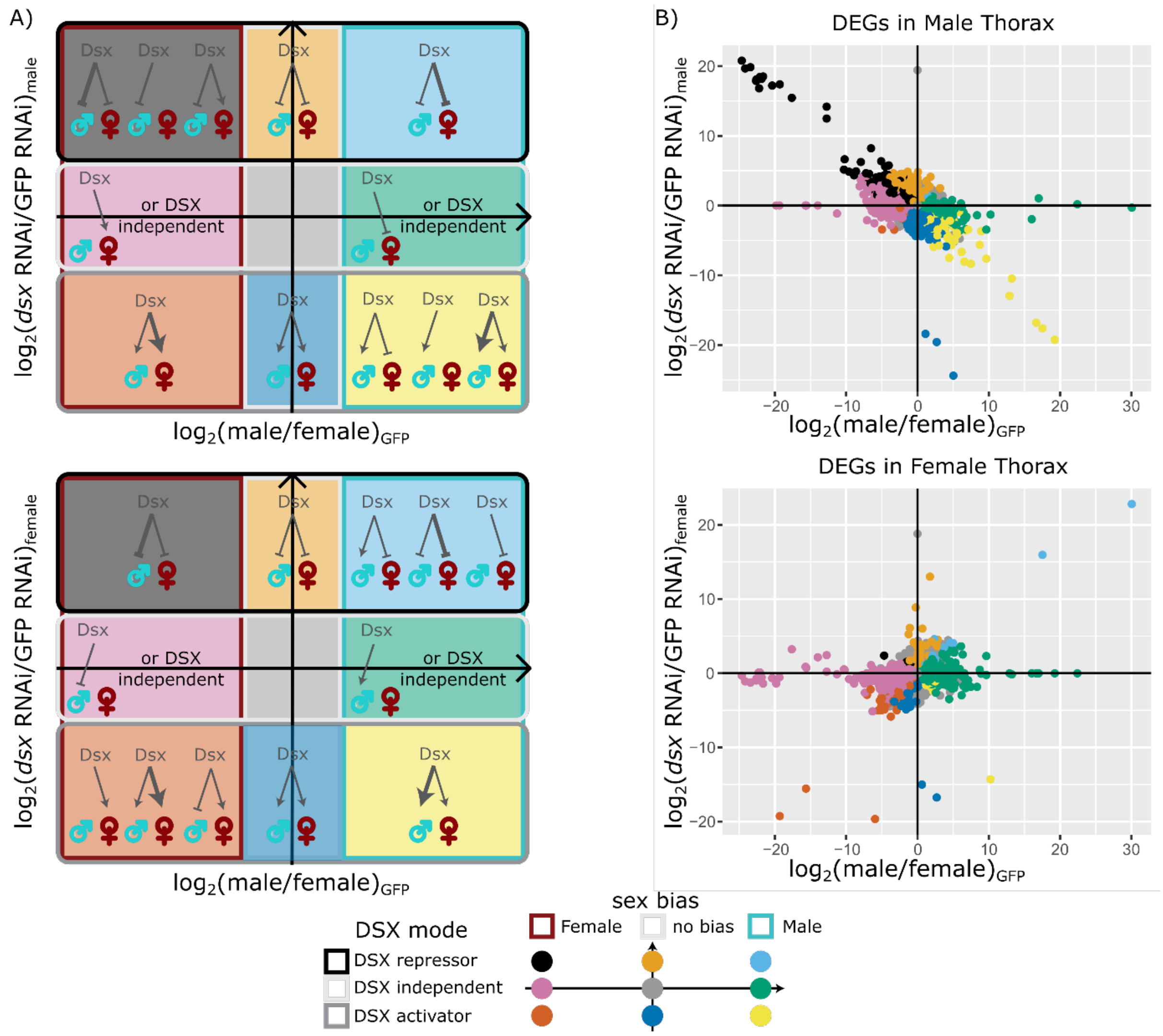


**Expanded View, Figure 1: DSX modes of gene expression regulation.**

**A)** Diagrams presenting the effect of *dsx* knockdown on gene expression in males (top) or females (bottom) as a function of sex-biased gene expression in the control condition (GFP RNAi) for a given time point. Different background colours define different interactions between sex bias and *dsx* knockdown effect on expression. For each interaction, possible modes of gene expression control by DSX are represented. Arrows represent expression activation, while blunt arrows represent repression. The arrow's thickness represents the DSX effect's hypothetical strength on gene expression, with a thick arrow indicating a strong effect and a thin arrow a moderate to low effect. Comparison between panel *dsx* knockdown effect in males (top) and females (bottom) allows discrimination between different DSX modes of gene expression control. For example, if a gene is female-biased and overexpressed in male *dsx* knockdown, it should appear in the top-left corner of the upper panel (black background). This regulation can be explained by three different DSX modes of regulation: DSX could downregulate the gene in males only, or it could upregulate the gene in females and downregulate it in males, or downregulate it in both males and females, yet with a stronger effect in males. Depending on the effect of *dsx* knockdown in females, the same gene would be found either on the lower panel's top, middle, or bottom part of the left side. The colour code representing the different combinations between *dsx* knockdown effects and sex bias is presented below the graphs. This colour code also applies to the graphs presented in panel "B".

**B)** DSX regulatory modes of action, defined as a function of interaction between *dsx* knockdown effect and sex bias, for each gene from thorax samples, as explained in "A". The Y-axis shows the effect of *dsx* knockdown on gene expression as fold change ($log_2$-transformed) between *dsx*- and *GFP*-RNAi male (top) or female (bottom) samples. For a given gene, DSX function is defined as activator when $log_2$FC(*dsx* RNAi/*GFP* RNAi)<0 and *Padj* < 0.05, and repressor when $log_2$FC(*dsx* RNAi/*GFP* RNAi)>0 and *Padj* < 0.05. The X-axis is identical for both graphs and presents fold change ($log_2$-transformed) between male and female control samples (*GFP* RNAi). We consider a gene to be male-biased when $log_2FC(male/female)_{GFP}>0$ and *Padj* < 0.01 and female-biased when $log_2FC(male/female)_{GFP}<0$ and *Padj* < 0.01. The colour code representing the different combinations between *dsx* knockdown effect and sex bias is presented below the graphs and corresponds to the colour code presented in panel "A".

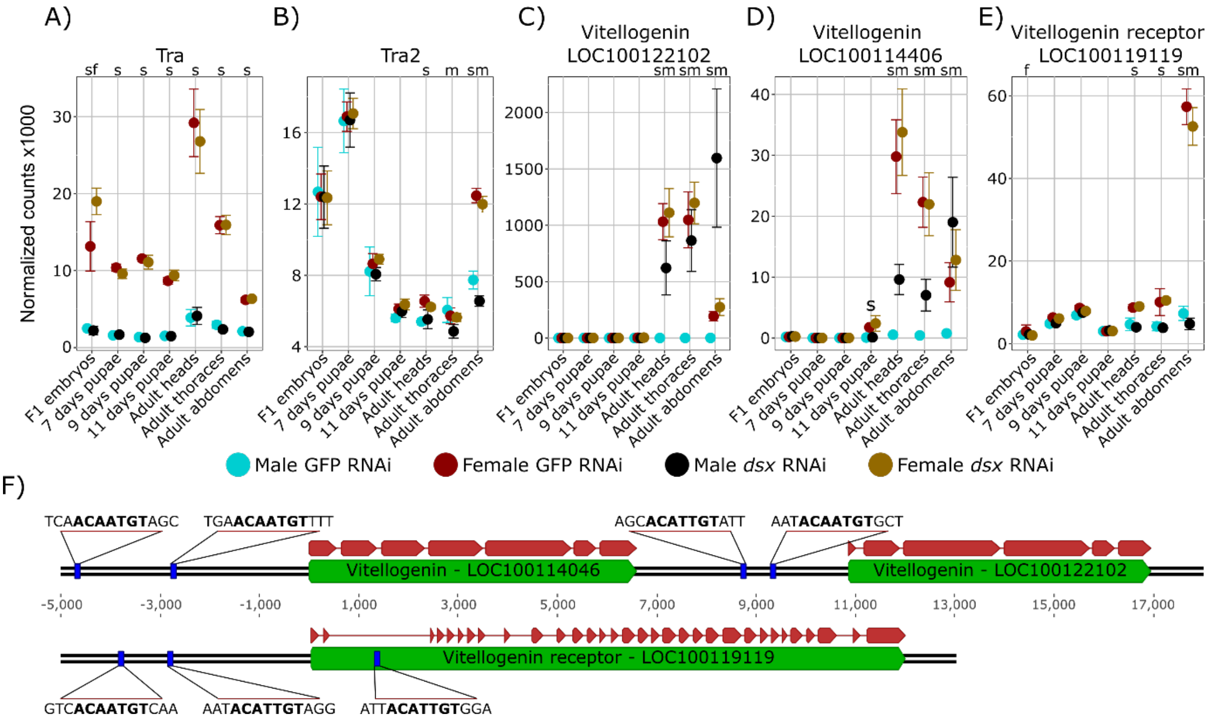


**Expanded View, Figure 2: DSX control on some notable genes expression**

**A-E)** Dot plots showing the expression level for a given gene in normalised counts after variance stabilisation transformation in DESeq2, plus and minus standard deviation. Above the graph is reported whether the differences between samples are statistically significant (adj. p value < 0.01): “s” = differences between *GFP*-RNAi males and females are statistically significant; “m”, differences between *dsx*-RNAi and *GFP*-RNAi males are statistically significant; “f”, differences between *dsx*-RNAi and *GFP*-RNAi females are statistically significant. *GFP*-RNAi male, *GFP*-RNAi female, *dsx*-RNAi male, and *dsx*-RNAi female are in turquoise, dark red, black, and brown, respectively.

**F)** Snapshot of *vitellogenin* (NC_045758.1 31113360..31136318) and *vitellogenin-receptor* (NC_045758.1 27482964..27500827) loci. Snapshots were taken using the Geneious (v2023.2.1) software and modified using Inkscape (v1.1.2). Green tracks represent the different genes annotated in the region, while blue tracks represent DSX binding sites identified in this study.

### Supplementary Information (Appendix)

**Supplementary table 1**: **Excel file containing the main results of our study regarding DSX role in sex-biased gene regulation.**

For each gene is reported the sex bias and the effects that *dsx* knockdown has on its expression in males and females, at each developmental stage and tissue we analysed. Moreover, we report for each gene the presence of a consensus DSX binding sequence of either 9 or 11 nucleotides. The information from these analyses was integrated to define a regulatory mode for DSX, as explained in **Supplementary Figure 3**; the regulatory mode is also reported in this table. Lastly, we include information regarding DSX regulation of the *Drosophila melanogaster* homolog according to Clough et al., (2014).

**Supplementary Table 2: Sex-bias expression and DSX mode of action for every gene annotated in the *Nasonia vitripennis* genome.**

DSX is classified as a repressor for a specific gene when the expression of this gene increased upon *dsx* knockdown in at least one sample from one sex, and it has never been observed to act as an activator. The classification is linked to the $DSX^M$ isoform when the effect is observed exclusively in males (DSX-M R), to the $DSX^F$ isoform when it is exclusive to females (DSX-F R), to $DSX^M$ or $DSX^F$ when it is found in both sexes but never in the same sample type (DSX-M or -F R), and to DSX double repressor when it is observed in both sexes within at least one similar sample type (DSX double R). DSX is classified as an activator for a specific gene when the expression of this gene decreased upon *dsx* knockdown in at least one sample from one sex, and it has never been observed to act as a repressor. The classification is linked to the $DSX^M$ isoform when the effect is observed exclusively in males (DSX-M A), to the $DSX^F$ isoform when it is exclusive to females (DSX-F A), to $DSX^M$ or $DSX^F$ when it is found in both sexes but never in the same sample type (DSX-M or -F A), and to DSX double activator when it is observed in both sexes within at least one similar sample type (DSX double A). DSX function is classified as changing when DSX has been observed acting both as an activator and repressor for a particular gene in two different samples within one sex. The function is linked to the $DSX^M$ isoform when the effect is observed exclusively in males (DSX-M C) and to the $DSX^F$ isoform when observed exclusively in females (DSX-F C). "Changing+Fix" identifies genes in which DSX effect changes within one sex but has a constant effect in the other sex. DSX function is classified as opposing when DSX has the opposite effect on a particular gene in the two sexes. The classification is divided into purely opposing when DSX activates a gene in one sex and represses the same gene in the other sex but in different developmental stages or tissues, and dual when a particular gene is activated in one sex and repressed in the other in at least one developmental stage or tissue. Finally, a mode of gene expression control is classified as independent of DSX when no effect of *dsx* knockdown could be observed on its expression.

| Mode | Mode (detailed) | Male | Female | Changing | No-Bias | total | % |
|---|---|---|---|---|---|---|---|
| Activator | DSX-M A | 1400 | 413 | 1385 | 118 | 3316 | 21 |
| | DSX-F A | 171 | 272 | 265 | 46 | 754 | 5 |
| | DSX-M or -F A | 29 | 99 | 70 | 19 | 217 | 1 |
| | DSX Double A | 6 | 5 | 10 | 3 | 24 | 0 |
| Repressor | DSX-M R | 460 | 1306 | 970 | 184 | 2920 | 19 |
| | DSX-F R | 173 | 98 | 145 | 69 | 485 | 3 |
| | DSX-M or -F R | 46 | 22 | 50 | 18 | 136 | 1 |
| | DSX Double R | 8 | 6 | 6 | 3 | 23 | 0 |
| Changing | DSX-M | 0 | 0 | 127 | 0 | 127 | 1 |
| | DSX-F | 0 | 0 | 2 | 0 | 2 | 0 |
| | Changing | 19 | 22 | 20 | 17 | 78 | 1 |
| | Changing+Fix | 9 | 3 | 10 | 1 | 23 | 0 |
| Opposing | Opposing | 47 | 64 | 54 | 14 | 179 | 1 |
| | Dual | 13 | 35 | 40 | 3 | 91 | 1 |
| Independent | | 1697 | 1167 | 1455 | 2777 | 7096 | 46 |
| | total | 4078 | 3512 | 4609 | 3272 | 15471 | |
| | % | 26 | 23 | 30 | 21 | | |

**Supplementary Table 3: Probability matrix of nucleotide occurrence in the DSX binding sequence.**

The DSX binding consensus sequence, [AT]TACA[TA]TGT[TA][TG][CT], was defined ex-novo with MEME (v5.1.0) on 903 single occurrences. In each row is reported the proportion of each nucleotide (A, C, G, and T) at a given position.

| **Position** | **[consensus]** | **A** | **C** | **G** | **T** |
|---|---|---|---|---|---|
| **-5** | [AT] | 0.661 | 0.075 | 0.014 | 0.25 |
| **-4** | [T] | 0.131 | 0.008 | 0.037 | 0.824 |
| **-3** | [A] | 0.999 | 0 | 0 | 0.001 |
| **-2** | [C] | 0.006 | 0.993 | 0.001 | 0 |
| **-1** | [A] | 0.867 | 0.003 | 0 | 0.13 |
| **0** | [AT] | 0.297 | 0.003 | 0.002 | 0.698 |
| **1** | [T] | 0.013 | 0.001 | 0.001 | 0.984 |
| **2** | [G] | 0 | 0.001 | 0.997 | 0.002 |
| **3** | [T] | 0.004 | 0 | 0.001 | 0.995 |
| **4** | [TA] | 0.345 | 0.034 | 0.007 | 0.614 |
| **5** | [TG] | 0.013 | 0.038 | 0.211 | 0.737 |
| **6** | [CT] | 0.161 | 0.494 | 0.134 | 0.211 |

**Supplementary Table 4: The ten most represented sequences in the DAP-seq dataset are also enriched in the genome.**

"p OBS/EXP" in DAP-seq is the binomial cumulative probability of observing these many or more sequences given how many NNNACAWTGTNNN sequences are present in DAP-seq dataset. "p OBS/EXP" in genome is the binomial cumulative probability of observing these many or more sequences compared to a random 13bp sequence in *Nasonia vitripennis* genome (NvPSR1).

| **Sequence** | **Occurrences in DAP-seq** | **Occurrences in genome** | **p OBS/EXP in DAP-seq** | **p OBS/EXP in genome** |
|---|---|---|---|---|
| **AATACAWTGTTTC** | 25 | 57 | <E-16 | <E-16 |
| **CATACAWTGTATT** | 24 | 55 | <E-16 | <E-16 |
| **TATACAWTGTATA** | 17 | 79 | <5E-12 | <E-16 |
| **GAAACAWTGTATA** | 16 | 36 | <E-16 | <1E-12 |
| **GAAACAWTGTATT** | 14 | 31 | <8E-15 | <1E-09 |
| **ATTACAWTGTTTC** | 12 | 26 | <3E-13 | <6E-07 |
| **AATACAWTGTATT** | 11 | 28 | <2E-11 | <6E-08 |
| **TATACAWTGTTGC** | 10 | 26 | <1E-10 | <6E-07 |
| **GCAACAWTGTATC** | 10 | 18 | <3E-12 | <0.002 |

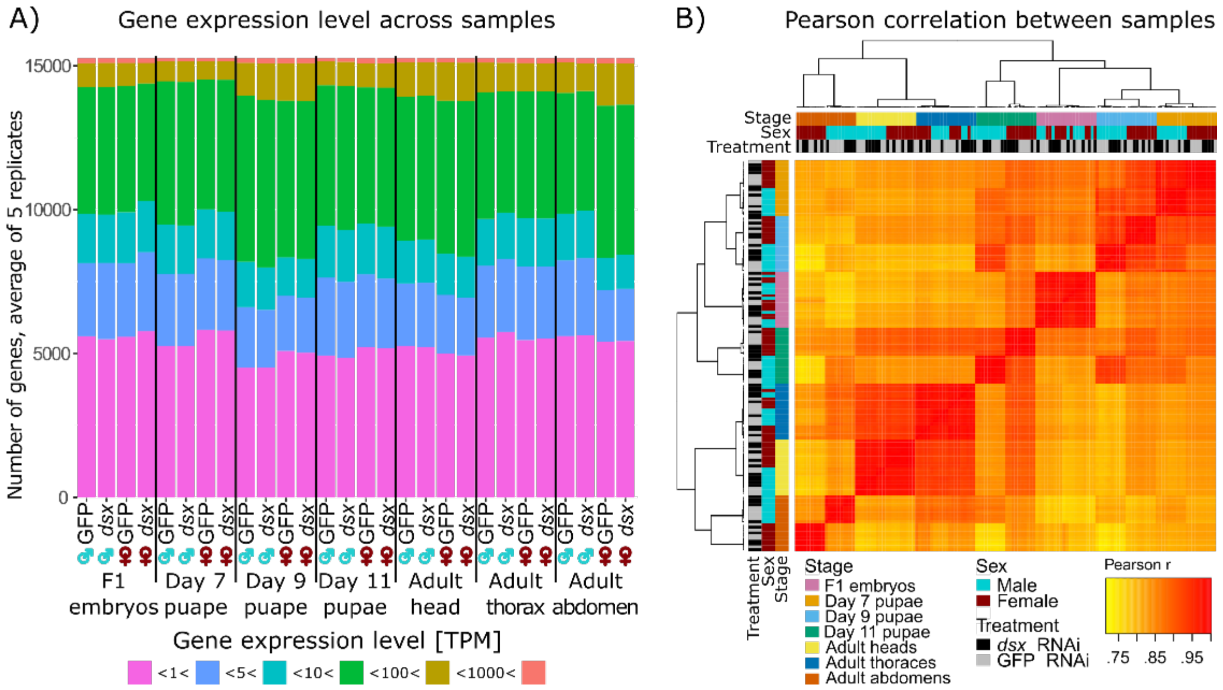


**Supplementary Figure 1: Quality Control report for all RNA-seq replicate samples generated in this study.** **A)** Bar plot showing the expression level of genes in transcripts per million reads (TPM) for every sample (average of 5 replicates). Genes were grouped into six categories based on their expression. All samples have approximately two-thirds of their genes expressed with TPM>1. Fucsia: genes with less than 1 TPM; blue: genes with more than 1 TPM and less than 5 TPM; aquamarine: genes with more than 5 TPM and less than 10 TPM; green: genes with more than 10 TPM and less than 100 TPM; gold: genes with more than 100 TPM and less than 1000 TPM; red: genes with more than 1000 TPM. **B)** Heatmap showing the Pearson correlation coefficient (r) between all samples. Samples were clustered by complete-linkage clustering with the R package "heatmap3" (version 1.1.7). Replicates have a high correlation (r>0.955). Samples cluster first by developmental stage or tissue and then by sex, except for embryos and thorax, where the sexes are mixed. Samples do not cluster based on *dsx*-RNAi or *GFP*-RNAi treatment.

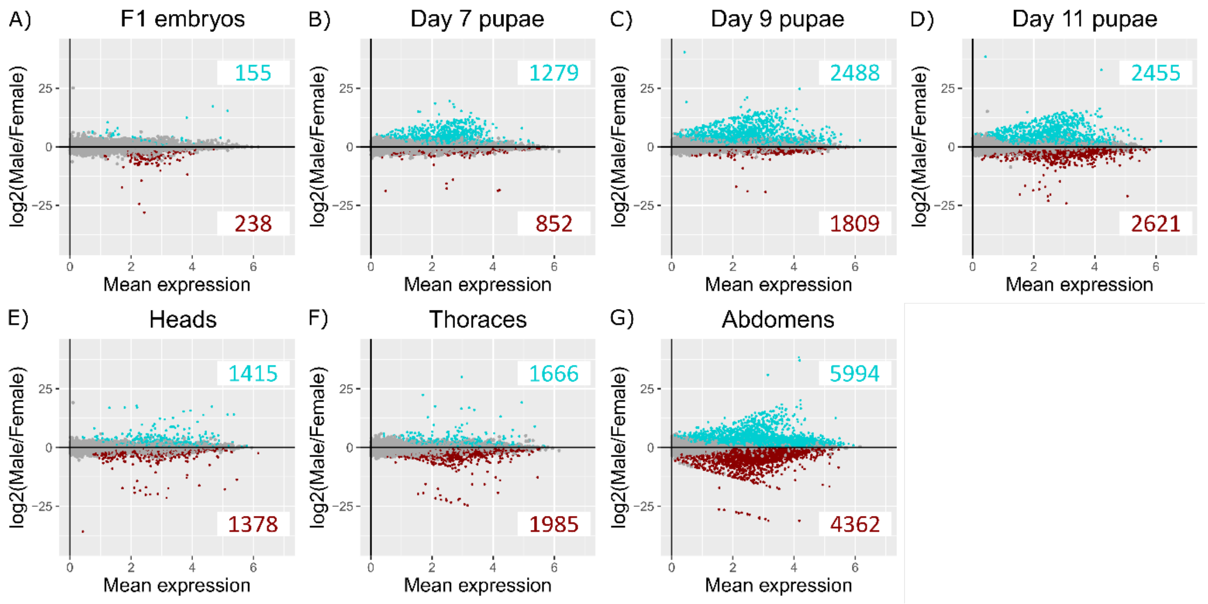


**Supplementary Figure 2:**

**A-G)** MA plots showing for each stage the fold-change ($log_2$-transformed) between *GFP* dsRNA injected control male and female samples as a function of the normalised average count between the two sexes ($log_{10}$-transformed) as calculated using DESeq2. The total number of genes significantly over-expressed or down-regulated are indicated in turquoise ($log_2FC>0$ and *P-adj*<0.01) and red ($log_2FC<0$ and *P-adj*<0.01) respectively, for each sample. Non-significantly deregulated genes in dark grey (*P-adj*>0.01).

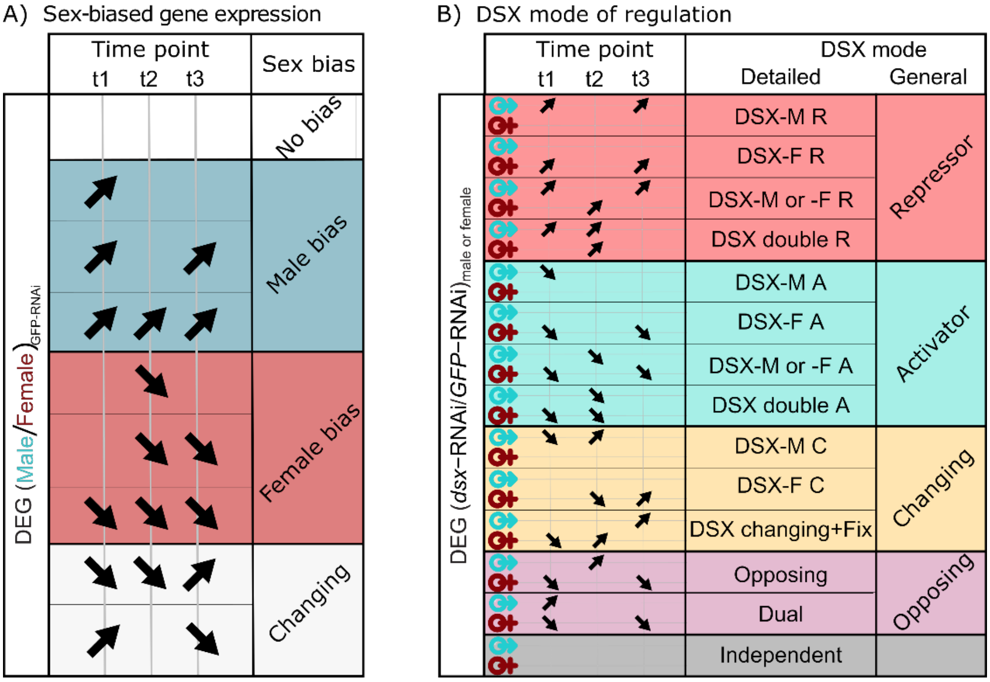


**Supplementary Figure 3: Defining the general mode of DSX regulation per gene across development.**

**A)** Example of how the general sex-biased expression per gene throughout development was determined. For clarity, an example with only three sample types is used. A gene has no sex bias when it is never identified as sex-biased in any sample. A gene is considered male-biased when it is identified as male-biased in at least one sample and is never female-biased. Similarly, a gene is considered female-biased when it is identified as female-biased in at least one sample and is never male-biased. A gene is considered changing in sex bias when it is identified as female-biased in at least one sample and male-biased in at least one other sample (grey).

**B)** Example of how the general DSX mode of regulation per gene throughout development was determined. For clarity, an example with only three sample types is used. DSX is classified as a repressor for a specific gene (red area) when it has been observed to repress this specific gene (represented by a downward-facing arrow) in at least one sample from one sex, and it has never been observed to act as an activator. The classification is linked to the DSX$^{M}$ isoform when the effect is observed exclusively in males (DSX-M R), to the DSX$^{F}$ isoform when it is exclusive to females (DSX-F R), to DSX$^{M}$ or DSX$^{F}$ when it is found in both sexes but never in the same sample type (DSX-M or -F R), and to DSX double repressor when it is observed in both sexes within at least one similar sample type (DSX double R).

DSX is classified as an activator for a specific gene (blue area) when it has been observed to activate this specific gene (represented by an upward-facing arrow) in at least one sample from one sex, and it has never been observed to act as a repressor. The classification is linked to the DSX$^{M}$ isoform when the effect is observed exclusively in males (DSX-M A), to the DSX$^{F}$ isoform when it is exclusive to females (DSX-F A), to DSX$^{M}$ or DSX$^{F}$ when it is found in both sexes but never in the same sample type (DSX-M or -F A), and to DSX double activator when it is observed in both sexes within at least one similar sample type (DSX double A). DSX function is classified as changing (yellow area) when DSX has been observed acting both as an activator (downward-facing arrow) and repressor (upward-facing arrow) for a particular gene in two different samples within one sex. The function is linked to the DSX$^{M}$ isoform when the effect is observed exclusively in males (DSX-M C) and to the DSX$^{F}$ isoform when observed exclusively in females (DSX-F C). "DSX changing+fix" identifies genes in which DSX effect changes within one sex but has a constant effect in the other sex.

DSX function is classified as opposing (lilac area) when DSX has the opposite effect on a particular gene in the two sexes. The classification is divided into purely opposing when DSX activates a gene in one sex (downward-facing arrow) and represses the same gene in the other sex (upward-facing arrow) but in different developmental stages or tissues, and dual when a particular gene is activated in one sex and repressed in the other in at least one developmental stage or tissue.

Finally, a mode of gene expression control is classified as independent (grey area) of DSX when no effect of *dsx* knockdown could be observed on its expression.

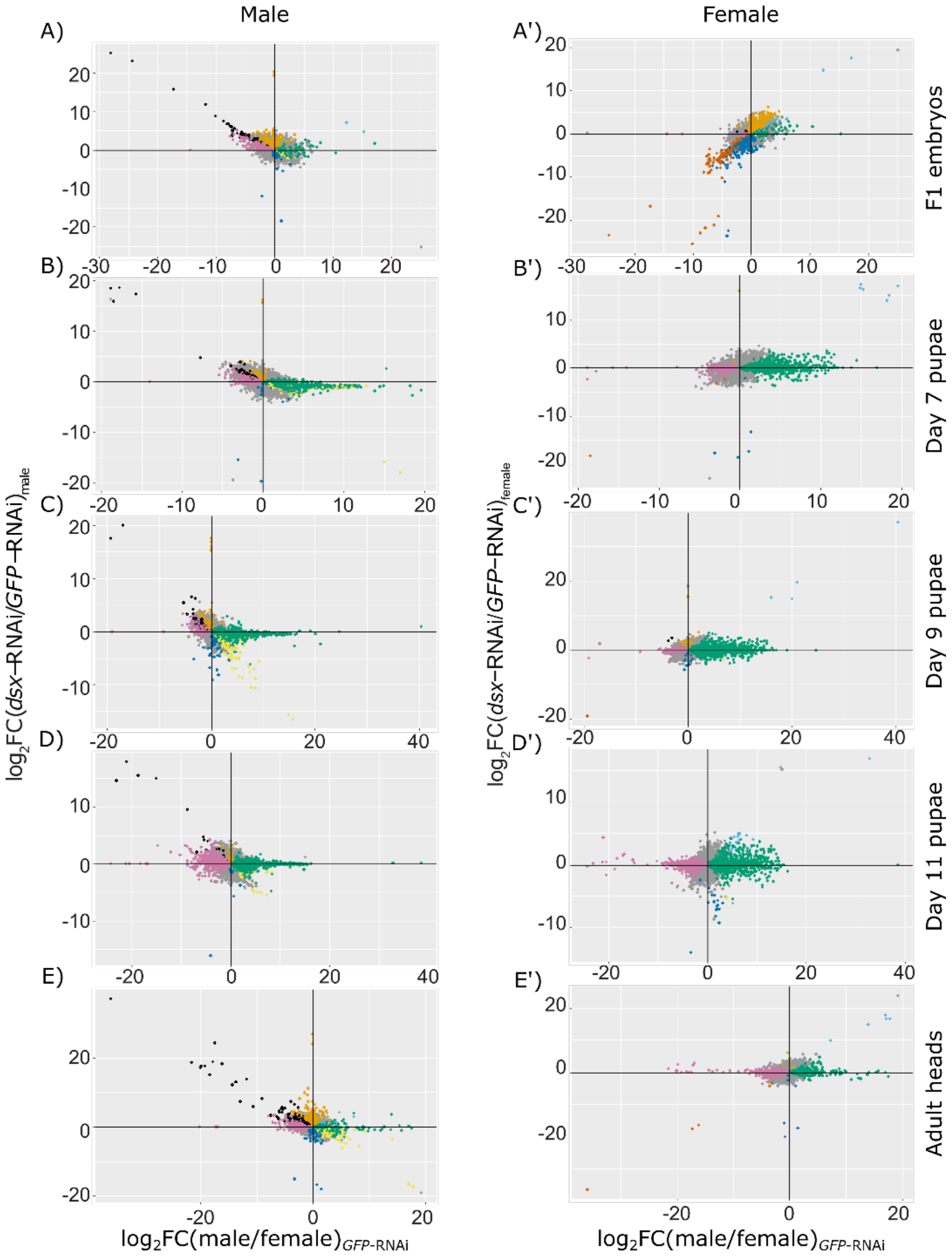
Male
Female
A)
A')
B)
B')
C)
C')
D)
D')
E)
E')
F1 embryos
Day 7 pupae
Day 9 pupae
Day 11 pupae
Adult heads
log2FC(dsx–RNAi/GFP–RNAi)male
log2FC(dsx–RNAi/GFP–RNAi)female
log2FC(male/female)GFP-RNAi
log2FC(male/female)GFP-RNAi

1286

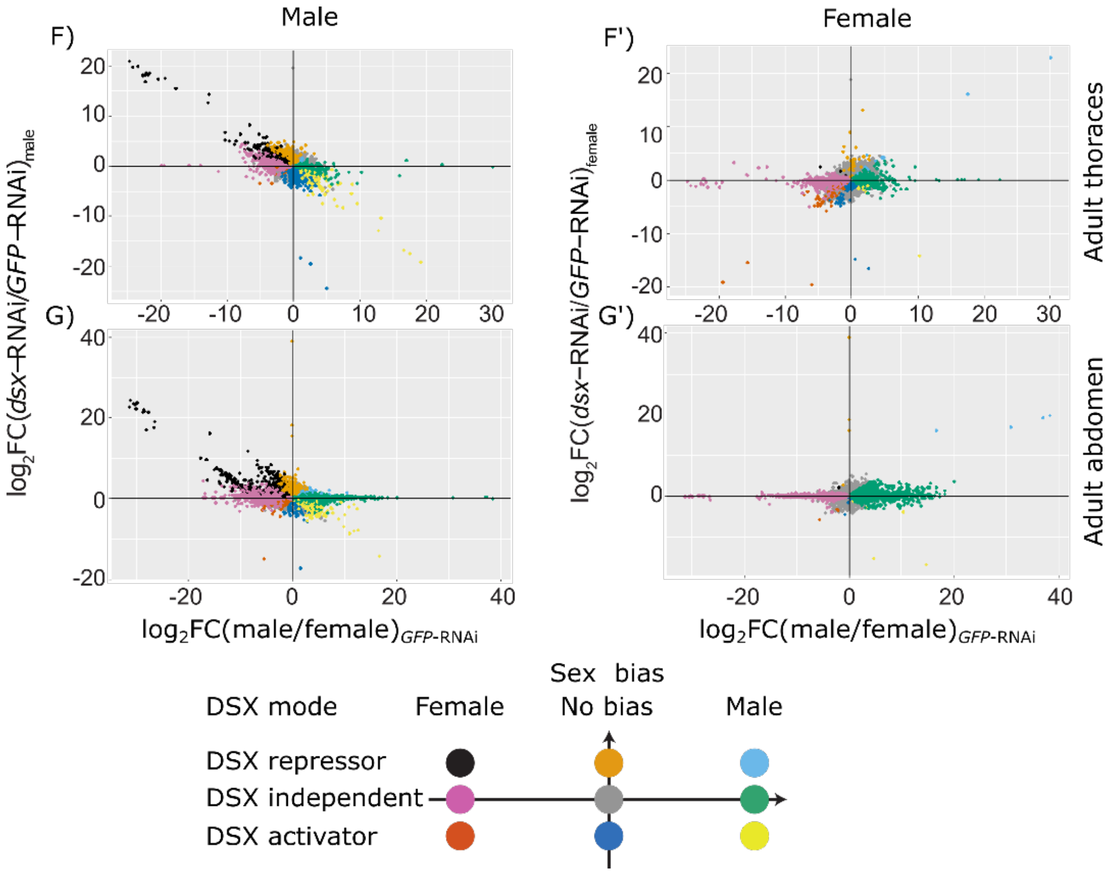


**Supplementary Figure 4: Identification of DSX modes of gene expression regulation for all sample types.** **A-G)** Dot plots showing the effect of *dsx*-RNAi as function of sex-biased expression for each sample type in males. **A'-G')** Dot plots showing the effect of *dsx*-RNAi as function of sex-biased expression for each sample type in females. For all graphs, the Y-axis shows the effect of *dsx*-RNAi on gene expression as fold change ($log_2$-transformed) between *dsx*-dsRNA and *GFP*-dsRNA injected samples. For a given gene, DSX function is defined as activator when $log_2$FC(*dsx*-RNAi/*GFP*-RNAi)<0 and *Padj* < 0.05, and repressor when $log_2$FC(*dsx*-RNAi/*GFP*-RNAi)>0 and *Padj* < 0.05. The X-axis is identical for both graphs and presents fold change ($log_2$-transformed) between *GFP* dsRNA injected control male and female samples. A gene is considered male-biased when $log_2FC(male/female)_{GFP}>0$ and *Padj* < 0.01 and female-biased when $log_2FC(male/female)_{GFP}<0$ and *Padj* < 0.01. The colour code representing the different combinations between DSX knockdown and sex bias is presented below the graphs and corresponds to the colour code presented in **EW Figure 1A** and **EW Figure 1B**.

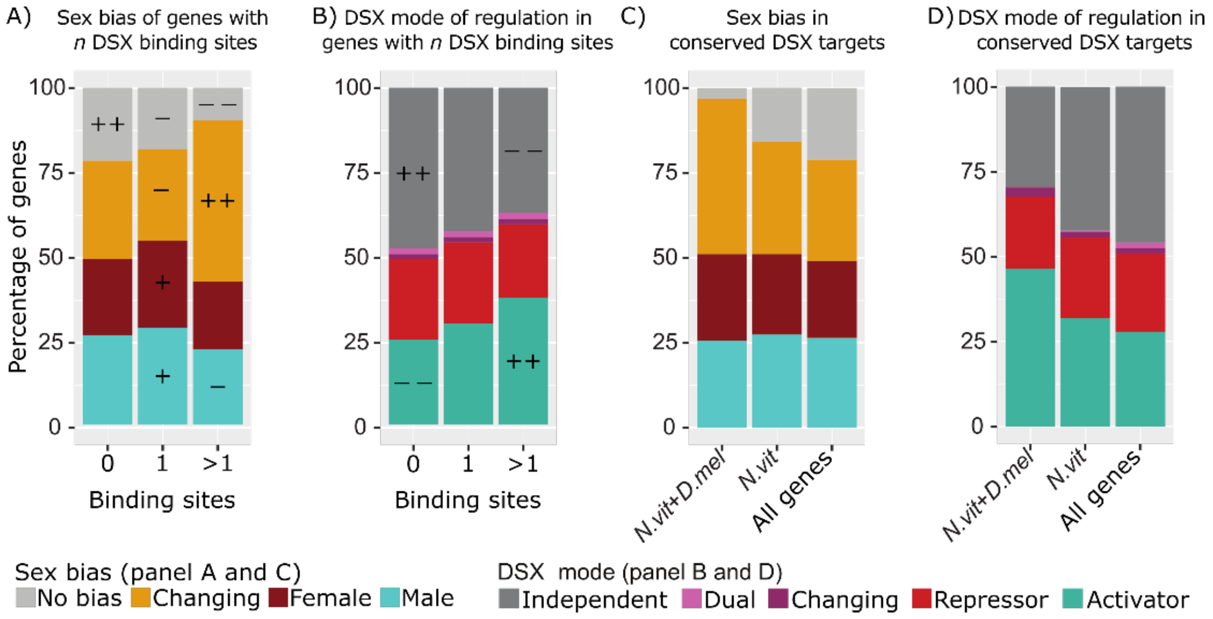


**Supplementary Figure 5: Repartition of genes based on sex-bias (A and C) or DSX mode of regulation (B and D) based on either number of DSX binding sites (A and B) or being a conserved DSX targets in *Nasonia vitripennis* and *Drosophila melanogaster* (C and D).**

**A)** Bar plot showing the proportion of genes per sex-bias classification as a function of DSX-binding site occurrence. The colour code is as follows: turquoise: male bias, dark red: female bias, ochre: changing sex bias, grey: no sex bias. Plusses and minuses represent category residuals relative to the $\chi^2$ comparison between expected and observed values: '++' = std. residuals > +4; '+' = std. residuals > +2; '-' = std. residuals < -2; '- -' = std. residuals < -4. Genes that change in sex bias are over-represented by genes with more than one binding site, while male and female-biased genes are over-represented in the group of genes with one binding site. Genes that are not biased are disproportionally represented in genes with zero binding sites. ($\chi^2$ (1, N = 15,274) = 233.0525, $p < 2.2e-16$).

**B)** Bar plot showing the DSX mode of gene expression control classification identified for a gene as a function of DSX binding sites occurrence. The colour code is as follows: red: repressor, aquamarine: activator, pink: dual, purple: changing, dark grey: independent of DSX control. Plusses and minuses represent category residuals relative to the $\chi^2$ comparison between expected and observed values: '++' = std. residuals > +4; '+' = std. residuals > +2; '-' = std. residuals < -2; '- -' = std. residuals < -4. Genes that are activated by DSX are disproportionally more represented in genes that have more than one DSX binding site. Genes that are not biased are disproportionally represented in genes with zero binding sites ($\chi^2$ (8, N = 15,274) = 222.6, $p < 2.2e-16$)

**C)** Bar plot showing the sex bias classification identified for each gene in *N. vitripennis* in different categories. "*N.vit+D.mel*" represent all potential targets identified in both *N. vitripennis* (this study) and in *D. melanogaster*, according to Clough et al., (2014); "*N.vit*" represent all potential targets identified in *N. vitripennis* (this study); "All genes" represents all genes in the *N. vitripennis* annotation. Definition of sex-bias categories are provided in **Supplementary Figure 3A**. Potential DSX targets that are shared between species are enriched in sex-biased genes in *N. vitripennis,* especially in the changing category.

**D)** Bar plot showing the DSX mode of gene expression control classification in *N. vitripennis* for a gene for different subgroups. "*N.vit+D.mel*" represent all potential targets identified in both *N. vitripennis* (this study) and in *D. melanogaster* according to Clough *et al.*, (2014); "*N.vit*" represent all potential targets identified in *N. vitripennis* (this study); "All genes" represents all genes in the *N. vitripennis* annotation. DSX mode categories are depicted as in **Supplementary Figure 3B**. Potential DSX targets conserved in both species are enriched in genes under DSX control in *N. vitripennis.*
